# *SongExplorer*: A deep learning workflow for discovery and segmentation of animal acoustic communication signals

**DOI:** 10.1101/2021.03.26.437280

**Authors:** Benjamin J. Arthur, Yun Ding, Medhini Sosale, Faduma Khalif, Elizabeth Kim, Peter Waddell, Srinivas C. Turaga, David L. Stern

## Abstract

Many animals produce distinct sounds or substrate-borne vibrations, but these signals have proved challenging to segment with automated algorithms. We have developed *SongExplorer*, a web-browser based interface wrapped around a deep-learning algorithm that supports an interactive workflow for (1) discovery of animal sounds, (2) manual annotation, (3) supervised training of a deep convolutional neural network, and (4) automated segmentation of recordings. Raw data can be explored by simultaneously examining song events, both individually and in the context of the entire recording, watching synced video, and listening to song. We provide a simple way to visualize many song events from large datasets within an interactive low-dimensional visualization, which facilitates detection and correction of incorrectly labelled song events. The machine learning model we implemented displays higher accuracy than existing heuristic algorithms and similar accuracy as two expert human annotators. We show that *SongExplorer* allows rapid detection of all song types from new species and of novel song types in previously well-studied species.

## Introduction

Animals produce diverse sounds (Kershenbaum et al., 2016), vibrations (Hill, 2006), and periodic electrical signals (Zakon et al., 2008) for many purposes, including as components of courtship, to sense their surroundings, and to localize prey. Quantitative study of these “sounds” is facilitated by automated segmentation. However, heuristic segmentation algorithms sometimes have low accuracy and fail to generalize across species (Arthur et al., 2013; Chesmore and Ohya, 2004; Coffey et al., 2019; Ivanenko et al., 2018; Koumura and Okanoya, 2016; LaRue et al., 2015; Parsons, 2001; Sattar et al., 2016). Song segmentation is particularly challenging for low signal-to-noise sounds, such as those produced by many insect species.

Deep neural network classifiers have been developed to study animal sounds (Chesmore and Ohya, 2004; Coffey et al., 2019; Ivanenko et al., 2018; Koumura and Okanoya, 2016; Parsons, 2001; Sattar et al., 2016; Steinfath et al., n.d.) and typically exhibit higher accuracy than heuristic algorithms. In addition, nonlinear dimensionality reduction techniques such as t-SNE (van der Maaten and Hinton, 2008) and UMAP (McInnes et al., 2018) have proven useful for discovering structure in animal sound datasets (Clemens et al., 2018). To facilitate adoption of deep networks and dimensionality reduction techniques for the analysis of animal sounds, we developed *SongExplorer* to support interactive workflows for discovery, annotation, and segmentation of animal sounds. We illustrate the utility of *SongExplorer* with examples from *Drosophila* courtship song (Ewing et al., 1968; Greenspan and Ferveur, 2000) because these low signal-to-noise songs have traditionally been challenging to segment (LaRue et al., 2015) and different species produce multiple distinct song types.

## Results

### *SongExplorer* provides a versatile interface for detecting and segmenting animal sounds

Each of the steps from discovery to automated segmentation of animal song events has traditionally required extensive manual investigation of songs and quantitative analysis, usually requiring use of multiple software tools. To democratize all steps of song analysis, we built a web browser-based program called *SongExplorer* that allows exploration of data, annotation of songs, and several analysis methods, including training of a deep learning neural network classifier, and quantitative assessment of classifier performance (Figure 1). The web interface presents users with three major domains in a single view (Figure 1A): the left side presents the data in multiple views; the middle section provides analysis “wizard” buttons, file handling, and parameter value windows; and the right side is a scrollable box containing extensive documentation and a tutorial.

**Figure 1.**
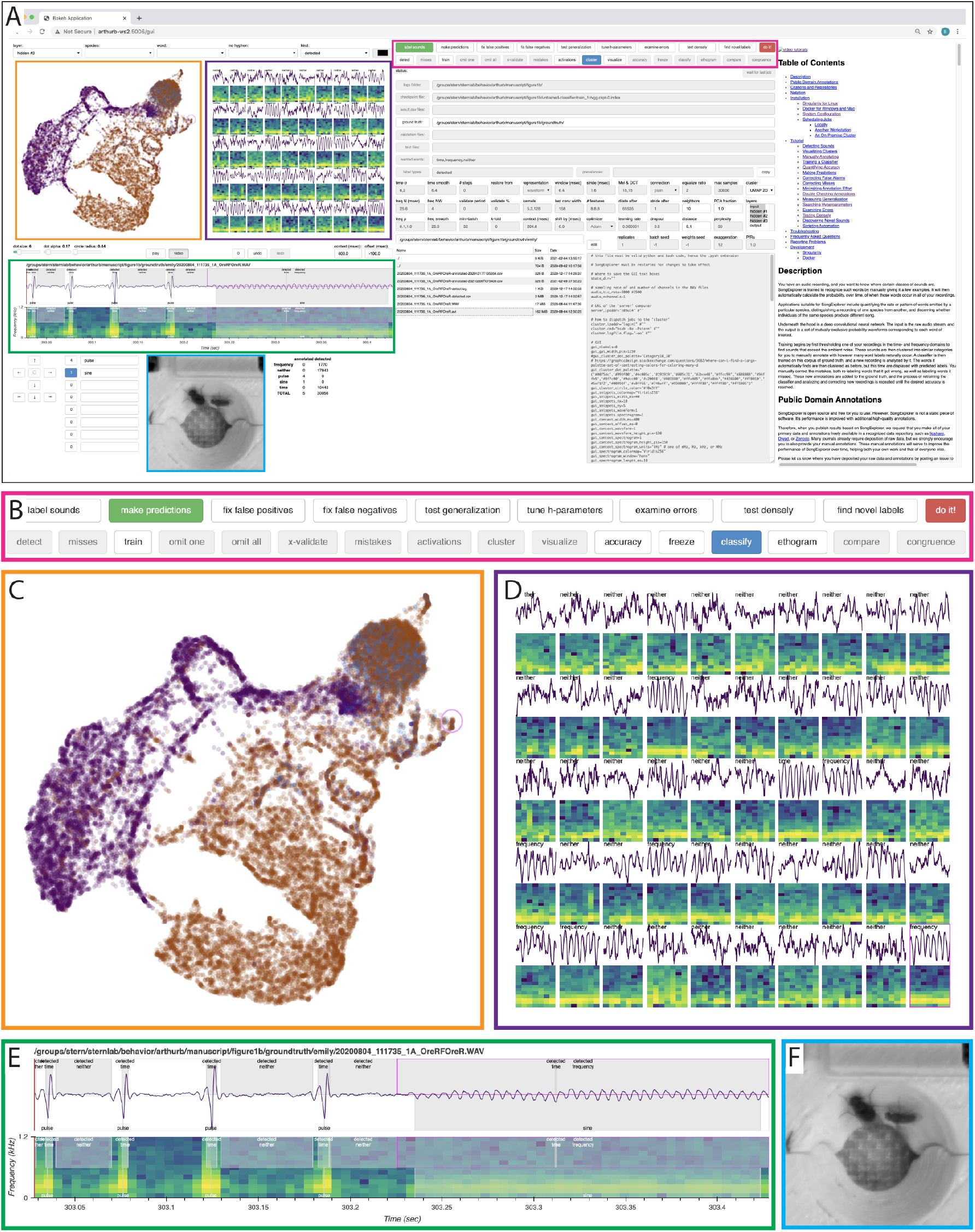
*SongExplorer* web browser interface (A) Screenshot of *SongExplorer* interface in a web browser window. Data and labelling tools are arrayed along the left side of wind. Analysis wizard buttons, file selectors, and parameter values are shown in the middle of the window. The right side of the window displays a detailed description of methods, including instructions for installation and data analysis. (B-F) Several regions of the interface are shown in more detail. The analysis wizard buttons (B) guide users, from left to right, through analysis pipelines and highlight only those windows that are available for each step of an analysis. The low-dimension feature embedding window (C) displays a tSNE or UMAP projection of sound events. These projections can be interactively navigated in two or three dimensions and different projections can be displayed for different song types, species, or layers of the machine learning model. A subset of events can be examined in detail by selecting events with an interactive circle selector (shown in pink here). The events, or a subset of events if there are more than 50 in the region, are shown along with spectrograms for the events (D). Clicking on one of these events results in the display of this event in the larger context of a complete recording (E). This view of the song can be navigated and played as an audio clip. Events can be labelled (or unlabeled) in this window. If video was captured at the same time as audio, then the synchronized video can be loaded and displayed in a window below the audio recording (F).

Quantitative study of animal sounds typically starts with supervised discovery of sounds. For species that produce loud and stereotyped sounds, like frogs and birds, it can be straightforward to identify individual types of songs. For other species, like many small insects, the initial step of identifying song types often requires examination of long recordings of audio and possibly video due to the sparse and quiet nature of their songs. Recent work has demonstrated that largely unsupervised clustering algorithms of song events can highlight distinct song types (Clemens et al., 2018). Therefore, to accelerate discovery, we provide methods for unsupervised clustering and visualization of song events.

The data view region on the left side of the browser window includes a box (Figure 1C) that displays data in a dimensionaly reduced form, either as a UMAP or tSNE representation, in either two or three dimensions. Samples of sound wave forms or frequency spectra can be projected into these reduced dimensionality spaces (Clemens et al., 2018). However, we have found that the hidden layer activations of a trained neural network provide more discrete representations of unique song types than do the original song events, which can facilitate identification of new song types (see later).

In *SongExplorer*, the reduced dimensionality space can be navigated rapidly with a modifiable “lens.” A sample of up to 50 sound events within the lens are represented in the adjacent window (Figure 1D). Raw sound traces are presented along with a spectrogram and a label indicating the song type. In the example shown in Figure 1, sounds were automatically detected (within *SongExplorer*) using thresholds either for high amplitude events (labelled “time”) or for events with a relatively strong signal in a subset of the spectrogram (labelled “frequency”). Clicking on each song event reveals the song event within a longer context of the recording in a separate window (Figure 1E). The context window (Figure 1E) can be navigated by zooming in or out and panning using buttons located below the context window. We have found that *Drosophila* song can sometimes be discriminated from other sounds most easily by listening to the song and watching associated video. Therefore, *SongExplorer* facilitates annotation by allowing the portion of the recording currently shown in Figure 1E to be played as audio and, if video data are available, for the video section for this region of the recording to be played. Individual song types can be named in boxes below the context window and labelled by using a computer mouse to double-click on events or to drag over ranges of continuous sounds. The number of annotated and automatically detected events are tracked below the context window.

The software is designed to encourage users to follow analysis pipelines that are enabled by “wizard” buttons (Figure 1B). For example, a user can explore a new dataset containing unlabeled data by selecting the “label sounds” wizard button, which enables five buttons that can be activated, from left to right, to perform the following steps: (1) automatically detect sounds above user-defined thresholds, (2) train a deep-learning classifier to recognize detected sound types, (3) calculate the classifier hidden-layer activations, (4) cluster these activation values, and (5) visualize the hidden layer activations using either UMAP or tSNE dimensionality reduction. Eight additional analysis pipelines are provided as wizards, and individual analysis steps can be implemented independently of the wizard buttons. We include methods to correct false positive predictions by a trained classifier and to iteratively retrain the classifier with corrected values. This kind of iterative classifier training has proven powerful for training classifiers to recognize visual behaviors (Berg et al., 2019) and served as an inspiration for our approach. We provide video tutorials (https://www.youtube.com/playlist?list=PLYXyXDkMwZip8x78RAyN6ee9NK42WBbKb) to guide users through these analysis pipelines.

### *SongExplorer’s* deep neural network displays higher accuracy than a heuristic classifier

*SongExplorer* includes a deep neural network classifier that can be trained to automate the annotation of animal sounds. The deep network is configurable, and defaults to a simple 3-layer time-domain convolutional neural network that classifies each sample in a 5000 Hz acoustic waveform using a 204.8 ms (1024 samples) window of context (Figure S1). We compared accuracy of this deep network versus a previously-described heuristic algorithm (Arthur et al., 2013) at classifying fly song. *D. melanogaster* males produce two distinctive types of courtship song, pulse song and sine song, and pulse song has traditionally been considered harder to classify than sine song. Kyriacou et al. (Kyriacou et al., 2017) performed a dense annotation of the pulse song events from 25 recordings of *D. melanogaster* courtship and a previously-described heuristic algorithm implemented on the same recordings, and with optimized parameters, displayed an F1 score (the harmonic mean of the precision and recall) of 87% (Stern et al., 2017).

To examine accuracy of the deep-learning neural network, we aligned pulse annotations to the nearest peak within five milliseconds and labelled all other points in time as “other” or “ambient” depending on whether a time-domain threshold was exceeded or not, respectively. We withheld five of the 25 recordings for validation and used the remaining 20 recordings to train the classifier. The classifier returned the probability that a pulse is centered on every time point in the recording (Figure 2A). To allow comparison with the heuristic algorithm, we implemented a discrete-valued ethogram by calculating a threshold based on the ratio of precision to recall; precision (also called the positive predictive value) is the fraction of true positives amongst all positives (true and false positives) and recall (sometimes called sensitivity) is the fraction of true positives detected amongst all real events (true positives plus false negatives). Using a precision to recall ratio of one (Figure S2), the trained network displayed an F1 of 94% (Figure 2B) for dense predictions made on the five withheld recordings, a considerable improvement over the heuristic algorithm. Within *SongExplorer*, users can select a lower or higher precision to recall ratio for thresholding to detect more of the relevant events (amongst more false positives) or mostly (but fewer) true positives, respectively. To explore the effect of sample size on classifier accuracy, we subsampled from the full dataset and found that 100 labelled pulse events produced accuracy very similar to the full dataset (Figure S3A). We also explored several parameters of the neural network and found that performance was largely insensitive to most hyperparameters (Figure S3B).

**Figure 2.**
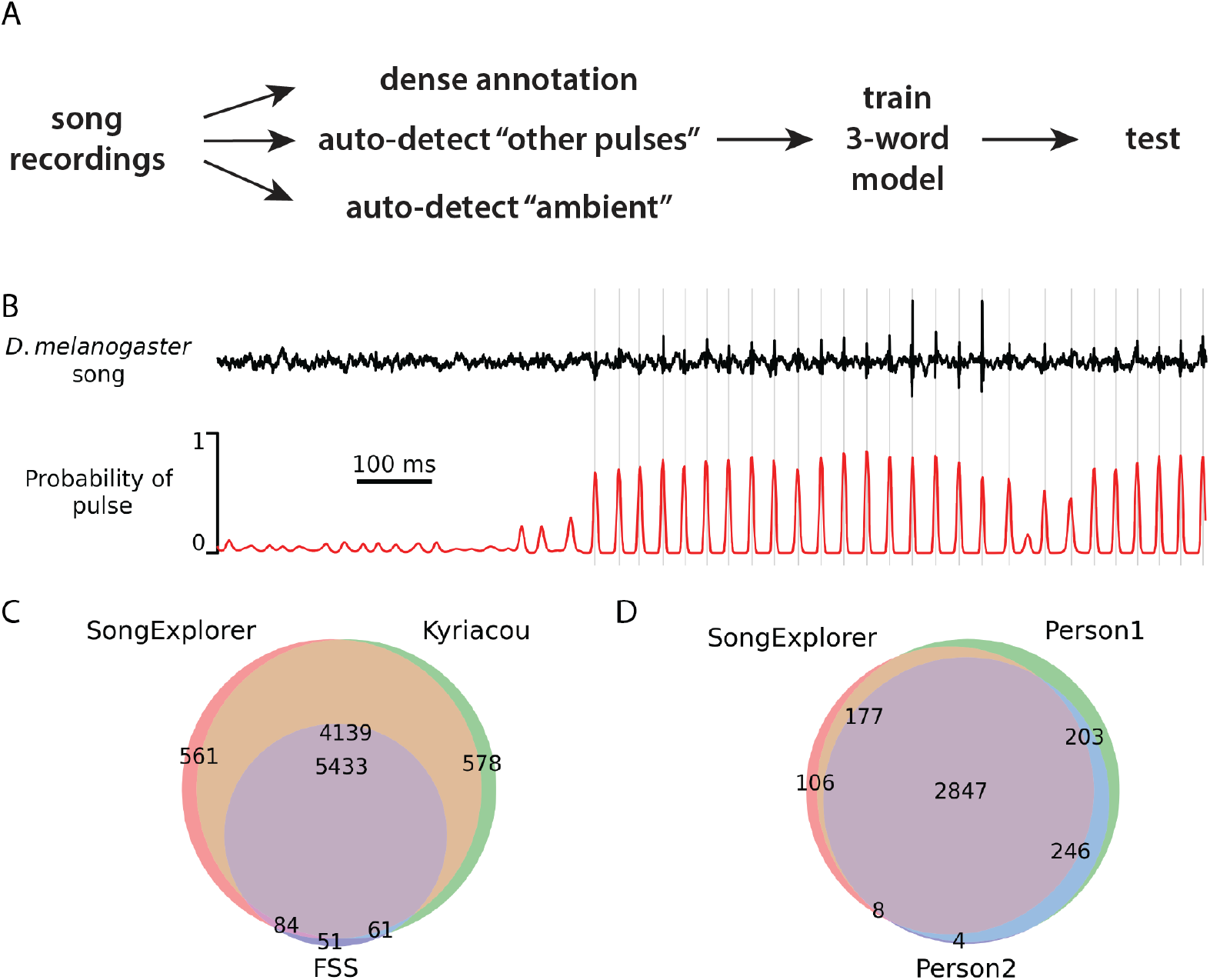
Performance of the neural network classifier to detect one kind of song event A – Outline of analysis pipeline employed to test accuracy of classifier to detect *D. melanogaster* pulse song. Dense annotations of pulse songs were combined with automated detection of other pulse-like sounds and ambient noise and a classifier was trained to recognize these three different kinds of sounds. Performance of the classifier was tested against dense human annotation. B – An example of a non-obvious train of pulse song from a *D. melanogaster* recording is shown above and the probability of a pulse event over time assigned by the classifier is shown below. Vertical grey lines indicate human annotated pulse events. C – To assess performance of the classifier, a dataset consisting of multiple recordings of many flies was annotated by an independent group that generated consensus labels of pulse events (“Kyriacou”) or segmented by our previously-developed classifier *FlySongSegmenter* (FSS). The neural network of *SongExplorer* was trained on 20 recordings and validated on the remaining five. 5433 pulses were labelled by all three methods (“Everyone”), 561 only by *SongExplorer* (*SongExplorer* False positives), 578 only by Kyriacou (False negatives) and 51 only by *FlySongSegmenter* (*FSS* false positives). A very large proportion of true pulses were not detected by *FlySongSegmenter* (4139; “not FSS”) but were detected by *SongExplorer*, illustrating the considerable improvement of *SongExplorer* over *FlySongSegmenter.* D – To determine how *SongExplorer*, trained on the Kyriacou consensus annotations, performs relative to individual humans, two authors with expertise in annotating fly song independently scored pulse events in the same random selections of fly song. *SongExplorer* and the two humans agreed on 2847 pulse events. *SongExplorer* displayed similar levels of disagreement on the remaining pulse events as the two humans. Subsequent manual investigation revealed that disagreements related to low-amplitude or isolated pulse events about which two humans could easily disagree.

### A trained classifier predicts pulse events approximately as well as an “average” human

To determine how well the trained neural-network classifier could generalize to predict pulse events in new recordings of *D. melanogaster* song, we made new recordings of *D. melanogaster* courtship song using chambers that were smaller than the chambers employed for the recordings described above. This is a challenging test case, because the noise characteristics differed systematically between the two sets of recordings (Figure S4). Two human experts independently performed dense annotations of pulse events for 23 randomly-chosen one-minute segments of these recordings without prior discussion of how they would label events. Different humans often disagree on annotation of low signal-to-noise events and, indeed, Person 1 labelled more events than Person 2. Both annotators and the classifier agreed on most pulse events (Figure 2C). Overall, the classifier and the two humans displayed similar levels of unlabeled events, suggesting that the classifier, even when trained on different sets of recordings, performed approximately as well as the “average” human.

### A common deep classifier architecture generalizes to many song types

Given the high accuracy of the classifier at detecting pulse song, we asked whether a classifier could be trained to accurately predict many song types across multiple species. We performed sparse annotations of all definable courtship song types from nine additional species, systematically labelled inter-pulse intervals between labelled pulse events, and trained a classifier to recognize each of the 37 song types (Figure 3A). Since these samples were labelled sparsely, we cannot examine accuracy as we did above for dense annotations. Instead, we examined the likelihood that an event was labeled correctly, given that there was an annotated song event at a particular time, and present the results as a confusion matrix (Figure 3B). The classifier assigned most events with greater than 90% accuracy, suggesting that the neural network architecture that we employed can be used to classify song from many *Drosophila* species.

**Figure 3.**
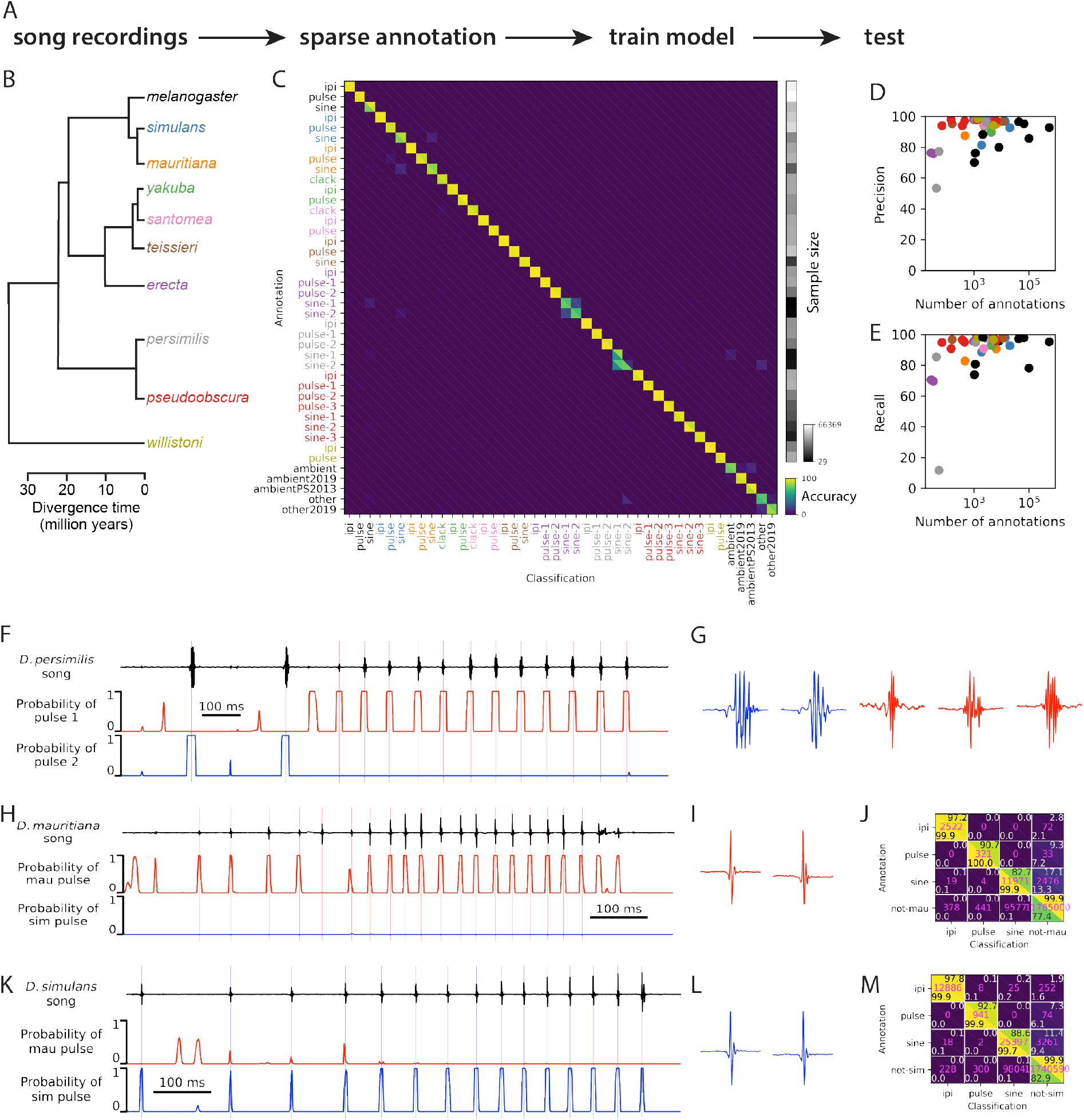
Multi-species song type classification by *SongExplorer* A – Analysis pipeline to assess ability of classifier to recognize songs from many *Drosophila* species. Song events from recordings of ten species were sparsely annotated and species-specific song-type labels were used to train the classifier. Classifier performance was tested by assessing the frequency with which the classifier correctly assigned a song type at manually annotated events. B – Phylogeny of the ten *Drosophila* species used in this analysis, color coded by the song types shown in axis labels and the performance statistics shown in panels C and D. C – Confusion matrix for 37 song types across ten *Drosophila* species color-coded by the species name given in B, plus “ambient” and non-song sounds (“other”). During manual annotation, we tentatively identified multiple types of similar songs for some species, which are indicated by numbers following the song types. Some of these alternative song types, such as *D. erecta* pulse types 1 and 2, are well discriminated by the classifier. Other types, such as *D. erecta* sine types 1 and 2 are not well discriminated, suggesting that they do not represent alternative song types. However, the sample sizes for alternative song types that are not well discriminated are low, suggesting that the failure to discriminate may have resulted from insufficient data for accurate training on these potentially different song types. Each colored square represents the fraction of time points that were annotated by humans as indicated by the row label and classified by the machine learning algorithm as indicated by the column label. The upper right triangles within each square sum to 100% within each row; the lower left sum within each column. Non-zero entries off the diagonal indicate false positive and false negatives in the lower left and upper right triangles, respectively. Each annotated time point was classified as the song type corresponding to the event with the highest probability. D, E — Precision (D) and recall (E) of the classifier for all song types plotted against the number of annotations for each song type. Performance tends to improve with increasing sample size for each song type. F – Example of classifier performance to discriminate two pulse types produced by *D. persimilis* males. A trace of approximately 1.5 sec of song is shown at top and the probabilities for pulse type 1 and 2 are shown below in red and blue, respectively. Vertical lines indicate human annotated pulse events color-coded by pulse type. G – Magnified views of the *D. persimilis* pulse types 1 and 2, shown in blue and red, respectively. H-M – The classifier discriminated similar pulse song events produced by two closely-related species, *D. mauritiana* and *D. simulans*, even though the differences are not obvious to humans. Song traces for *D. mauritiana* (top of G) and *D. simulans* (top of J) are shown with the classifier probabilities shown below each trace. Magnified views of example pulses from *D. mauritiana* (H) and *D. simulans* (K) do not reveal obvious differences between the pulses. Confusion matrices for *D. mauritiana* (I) and *D. simulans* (L) reveal that the classifier classifies pulse events to the correct species with greater than 90% accuracy. Vertical lines in (H) and (K) indicate human annotated pulse events color-coded by species, red for *D. mauritiana* and blue for *D. simulans*.

The classifier discriminated with high accuracy similar song types within a species and very similar song types from closely related species. For example, *D. persimilis* produces two different pulse types (Figure 3D-E) and the classifier accurately discriminated between these subtypes (Figure 3B). Surprisingly, the classifier also accurately distinguished between the very similar pulse events of the sister species *D. simulans* and *D. mauritiana* recorded on the same set of microphones, which, in our experience, cannot be discriminated by humans (Figure 4F-K). The classifier employs a context window of 204.8 ms surrounding each event, and the classifier may therefore have used information about the diverged inter-pulse intervals between species to discriminate between these pulse types.

**Figure 4.**
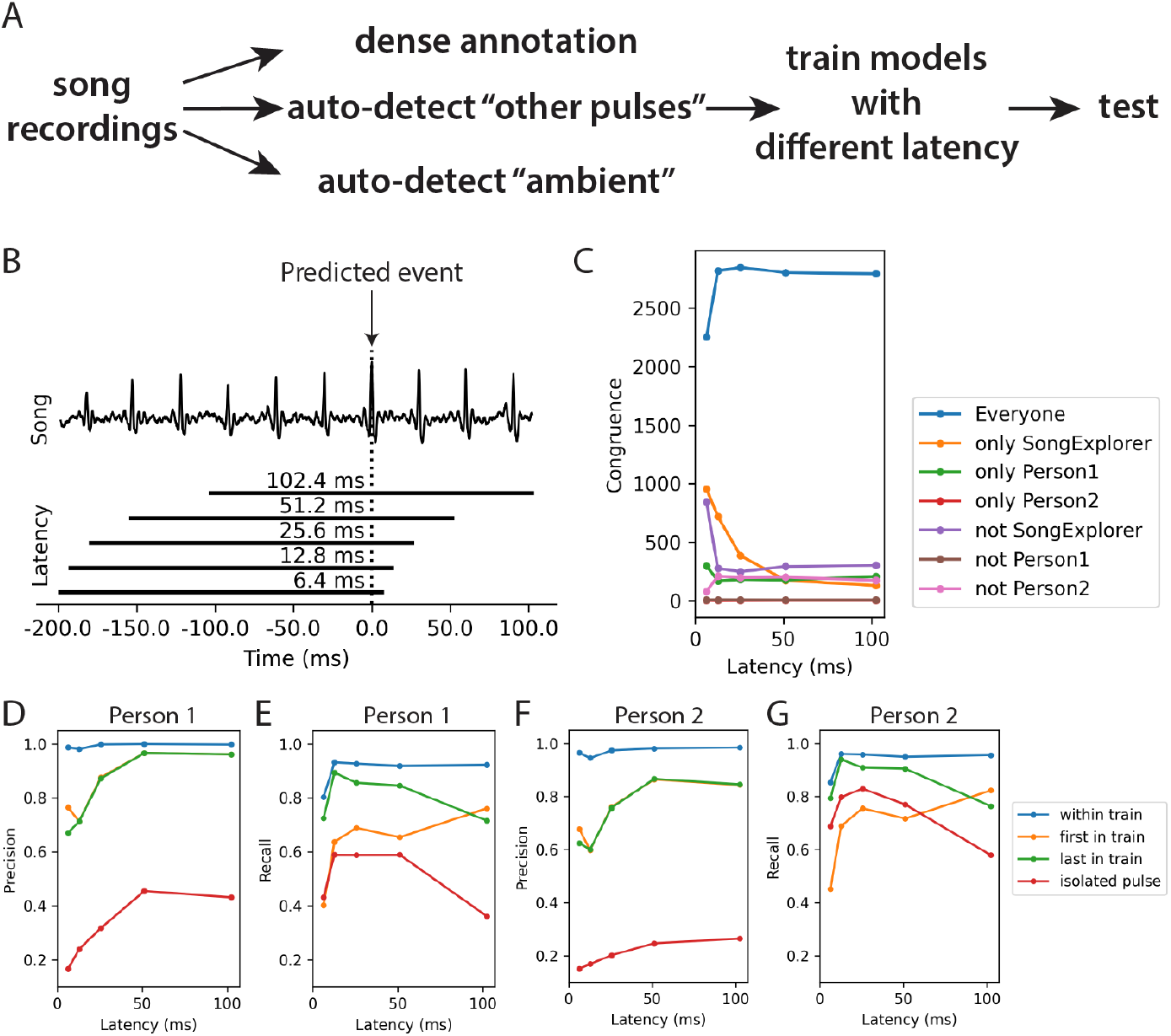
*SongExplorer* allows fast prediction, which will facilitate closed-loop applications. A – Outline of the analysis pipeline, which differs from the analysis pipeline shown in Figure 2A only by the use of multiple context windows, representing different latencies relative to the predicted event, for training. B – Illustration of how the context window was shifted to test ability of neural network to predict event at the focal position, indicated by dotted vertical line. Example shows song trace containing ten pulse events. The context window, which is normally centered on the predicted event, was shifted earlier in time to test the predictive ability of the classifier when the latency between the event and the prediction is shifted from 102.4 ms to 6.4 ms. C – Performance of the classifier with different latencies was measured as congruence amongst human annotators and the deep learning network. Congruence amongst “Everyone” remained high for all latency durations except for 6.4 ms. D-G – Classifier performance using different latencies relative to manual annotation by two humans. Performance was considerably lower for isolated pulses relative to pulses within trains. Classifier performance for the first and last pulse of each train were also lower than for pulses within trains.

Some song types were not discriminated well, such as *D. erecta* sine 1 vs sine 2 and *D. persimilis* sine 1 vs sine 2. In both cases, the alternative labels were assigned during manual annotation prior to the availability of *SongExplorer. Post hoc* examination of songs within the *SongExplorer* interface revealed that sine song is rare in both species and there is no compelling evidence for multiple sine types, suggesting that the classifier correctly failed to discriminate between sine song subtypes in these species because the species do not produce multiple subtypes. Alternatively, it is possible that the classifier failed to discriminate multiple sine song types because these songs are rare. However, we found weak dependence of classifier accuracy and precision on song event sample size (Figure 3C) and sample sizes above 100 had similar accuracies.

### A trained classifier can provide rapid prediction of song events for closed-loop assays

Rapid prediction of song could be valuable for closed-loop experiments. We therefore examined whether our classifier could be implemented, in principle, in closed-loop scenarios. We examined classifier accuracy when the 204.8 ms context window (the position of the sampled window relative to the target song event) was shifted earlier in time (Figure 4B). We call the time between the predicted event and the end of the context window the latency. This latency corresponds to “information latency” and is a lower bound on the true latency, which will be higher and depend on the efficiency of the software and hardware implementation, which we do not consider here. The classifier had precision of 88.6% and recall of 76.6%, on average, in predicting *D. melanogaster* pulse events with a latency as low as 12.8 ms (Figure 4B-F), compared with precision and recall of 91.5% and 89%, respectively, with a latency of 102.4 ms. Accuracy was lower for isolated pulses and the first pulse in a pulse train (Figure 4D-G), suggesting that the classifier may have used past pulses in a pulse train to improve the accuracy of detecting future pulses. Thus, in principle, this neural network classifier can identify many song events on the order of fast neuronal spiking in *Drosophila* (~200Hz).

### The learned feature representations of the neural network exhibit latent structure about song types and allow efficient discovery of new songs

Investigators studying animal sounds typically spend considerable time listening to recordings and examining song traces and video to identify and categorize sounds. This work is not only time consuming and tedious, but it is also subjective and can frustrate identification of rare sounds. We therefore sought a method to rapidly identify both common and rare sounds. We found that the activities of the hidden-layer neurons of a trained network exhibit considerable structure about song types (Figure 5) and allow rapid identification of novel song types. We illustrate two modes of this discovery approach.

**Figure 5.**
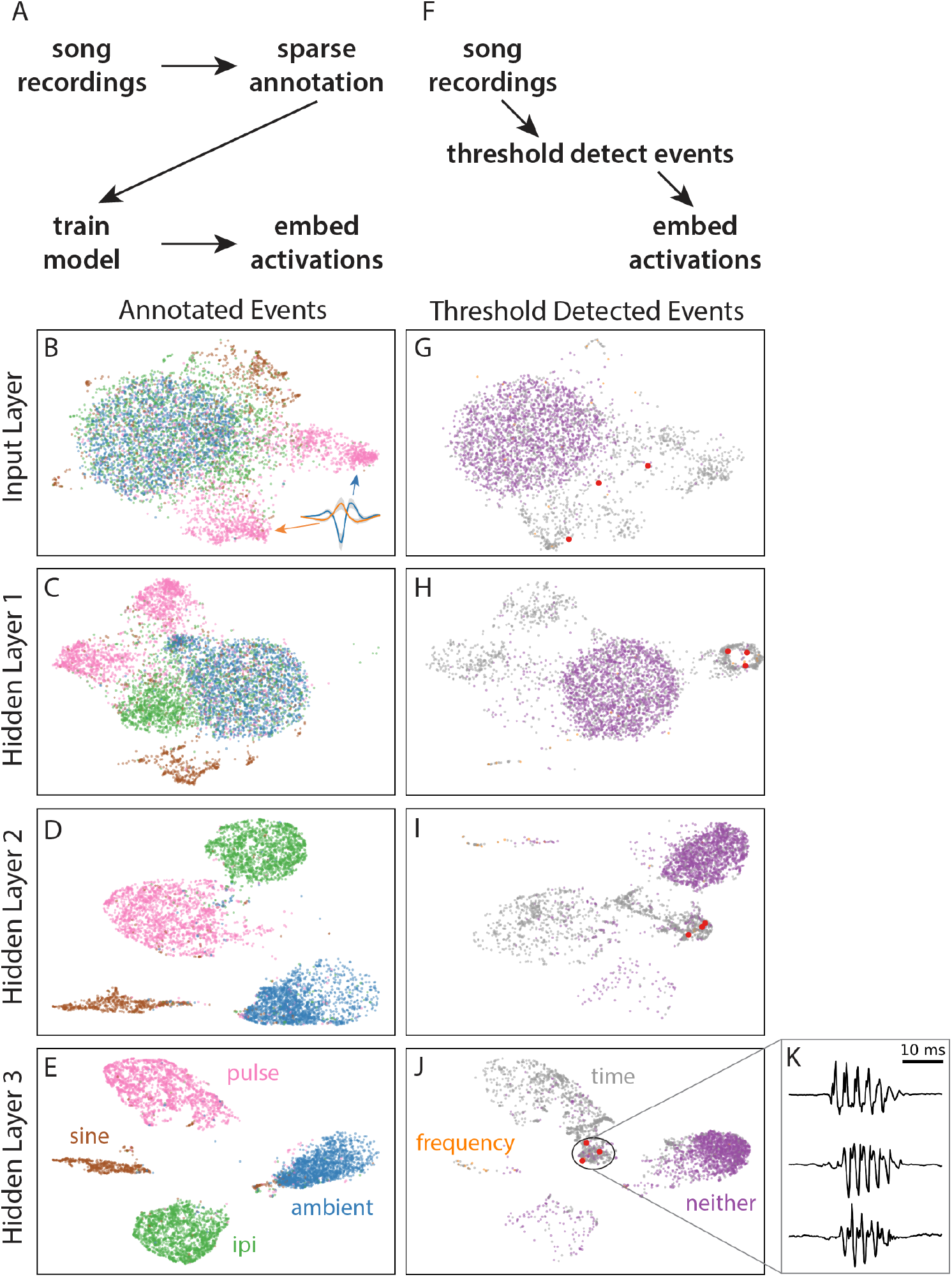
Dimensionality reduction reveals structure of hidden layers activation and facilitates rapid identification of new song types. A – Analysis pipeline to illustrate how visualization of hidden layer activations reveals relatively discrete structure of single song types. Songs that were sparsely annotated for pulse, sine, inter-pulse interval and ambient sounds were used to train a classifier. Principle component analysis values of the the input and hidden layer activations were projected into UMAP space. B-E – UMAP projections of input (B) and hidden layer (C-E) activations for *D. melanogaster* song. For input layer (B), pulse events form two loose clusters and sine song forms one loose cluster and all IPI events overlap with ambient sound. The two pulse event clusters represent the two phases of pulse song present in the data, which is an artifact of the position of the fly relative to the microphone at different times in the recording. At increasingly deeper layers in the model, the activations become increasingly differentiated (C-E), and clearly separate out most pulse, sine, IPI, and ambient events by hidden layer 3 (E). F – Analysis pipeline illustrates how new song types can be rapidly identified by clustering hidden layer activations. Sound events passing a simple amplitude or power spectrum threshold were projected through the model trained on annotated song (A-E) and the input layer and hidden-layer activations were projected into UMAP space. G-K – One strong cluster of thresholded events was obvious in all hidden layers (H-J), but not obvious in the input layer (G). Most of these events corresponded to a novel song type, which has recently been reported to be the female copulation song. Three of these copulation song events are illustrated (K) and their locations in the input and hidden layers are indicated by three larger red dots.

First, we trained a network on manually labelled male *Drosophila melanogaster* pulse and sine song. We also included automatically generated labels for inter-pulse interval and ambient noise samples in the training. We visualized the patterns of hidden layer neural activation in UMAP space (Figure 5A-D). These representations revealed that the input layer showed some structure, with pulse song mainly occupying two domains of the UMAP space and sine song a third domain (Figure 5B). However, points were distributed diffusely and multiple song types were intermingled. The two domains of pulse song in the input layer reflect the two phases of pulse song resulting from different positions of the fly relative to the directional microphone (Figure 5B inset). Through successive layers of the network, however, we noticed that each of the song types coalesced into nearly distinct clouds corresponding to the trained label classes of pulse song, sine song, inter-pulse interval, and ambient noise (Figure 5C-E). The network apparently correctly learned that the positive and negative deflecting phases of pulse song were an artifact and not distinguishable features of pulse song.

To explore whether other types of songs might have been missed in *D. melanogaster* recordings, we selected a different set of recordings, which had not been manually annotated, and detected all sounds using time- and frequency-domain thresholds. These sounds were projected through the neural network and the neural activation values were embedded in a common UMAP space defined by annotated and detected song events (Figure 5F). This embedding revealed a new density of points that was obvious in the hidden layers (Figure 5F-H), but not in the input layer (Figure 5E). Manual examination of these events (a subset of which are highlighted in red in Figure 5G-J) revealed that they are examples of a recently described female copulation song (Kerwin et al., 2020). Thus, this approach to discovering animal sounds allows rapid discovery of known and previously unknown song types in large audio datasets.

To further explore the utility of this kind of approach to song discovery, we trained a classifier to distinguish samples from 21 species (Figure 6A), providing as training data only sound events detected by time- and frequency-domain thresholding. Although the classifier was trained only to recognize the species of origin for a song event, the hidden layer activations contained considerable latent structure that facilitated discovery of song types (Figure 6B-D). That is, without providing *any* manually labelled training data, *SongExplorer* revealed the multiple song types produced by each species (Figure 6E-F). This allowed discovery of new song types in species of the *Drosophila nasuta* species group. One previous paper has reported song types from some of these species (Hongguang et al., 1997). Notably, we identified multiple song types in several species that were not identified in the earlier study (Figure S6). We estimate that using *SongExplorer* we discovered many or all song types for each species within approximately 20 minutes per species of exploring songs.

**Figure 6.**
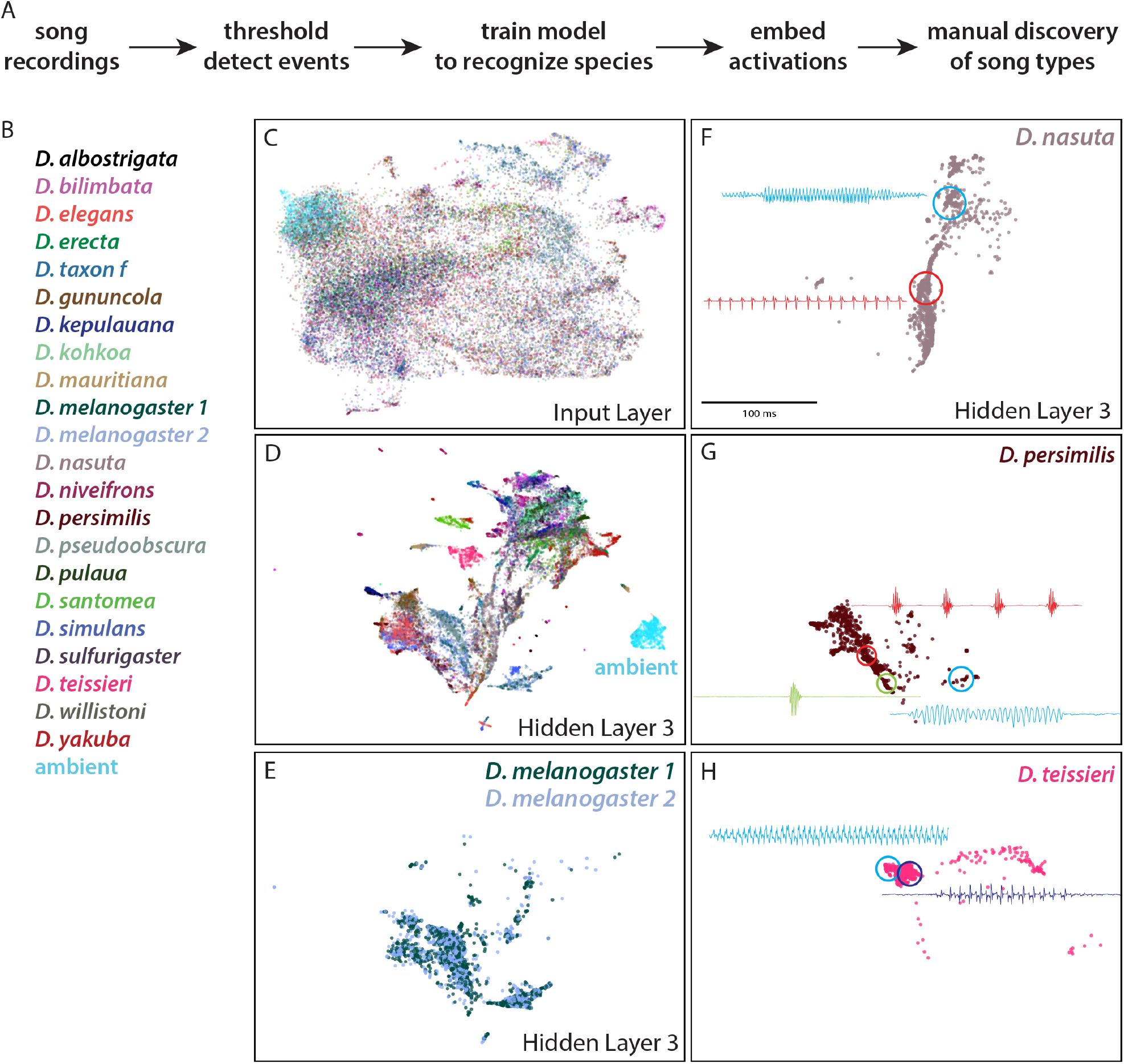
A classifier trained to recognize species sounds, rather than specific events, facilitates *de novo* discovery of song types. A – The deep learning classifier was trained to recognize sounds automatically detected with time- and frequency-domain thresholds sampled from multiple species. The classifier was trained to recognize the species, not individual song types. Data represented in UMAP space were manually examined to identify previously discovered and new song types. B – The 21 species used in the analysis. Two *D. melanogaster* recordings were labelled separately as a negative control. C,D – UMAP projections of the input layer (C), which contained little structure, and hidden layer 3 (D), which exhibited extensive structure. The detected sound events from many species occupied distinct domains in the UMAP projection of hidden layer 3. E – UMAP projections of hidden layer 3 for detected events from the two *D. melanogaster* recordings shows that that the classifier was unable to differentiate two sets of recordings from *D. melanogaster*. F-H — UMAP projections of *D. teissieri*, *D. nasuta*, and *D. persimilis* illustrate that songs from different species occupy different locations within UMAP space and facilitated discovery novel song types.

## DISCUSSION

*SongExplorer* provides the first interactive graphical interface to allow exploration, discovery, and segmentation of animal sounds using deep learning tools. While we have characterized *SongExplorer* and a particular instantiation of a neural network model using *Drosophila* courtship song, many kinds of animal sounds can be accurately classified using deep learning models (Chesmore and Ohya, 2004; Coffey et al., 2019; Ivanenko et al., 2018; Koumura and Okanoya, 2016; Parsons, 2001; Sattar et al., 2016; Steinfath et al., n.d.). The neural network classifier accuracy is only weakly dependent on sample size and the classifier attained greater than 90% accuracy at detecting *D. melanogaster* pulse song with just 100 labelled events.

Many researchers will be interested in generating discrete ethograms from the probabilistic output generated by *SongExplorer*. There are many ways to discretize the *SongExplorer* output and for this study we employed a threshold derived from the precision-recall curve. It is also possible to employ heuristic thresholds, or combinations of heuristics to filter *SongExplorer* output for any particular purpose.

*SongExplorer* provides an intuitive interface for efficient discovery of new song types, segmentation of large song datasets, and reannotation of songs to correct false negative and false positive labels. In contrast to the size of deep learning models used in computer vision, our deep network for acoustic signals is light-weight and runs acceptably on a CPU without the need for specialized GPU hardware. We therefore anticipate that *SongExplorer* may be valuable to a wide range of biologists studying animal sounds.

## Materials and Methods

### *SongExplorer* software program

The *SongExplorer* user-interface is web browser-based and is implemented in Python using the Bokeh library (https://bokeh.org/). It is cross-platform and has been tested to run on Mac OS X, Windows, and Linux computers. The deep learning components use Keras and Tensorflow and benefit modestly from access to a CUDA-compatible Nvidia GPU.

### Software availability

We provide the source code as well as Singularity and Docker containers of *SongExplorer* for Linux, Microsoft Windows, and Apple Macintosh platforms (https://github.com/JaneliaSciComp/SongExplorer). Extensive tutorials on using *SongExplorer* are available on YouTube (https://www.youtube.com/playlist?list=PLYXyXDkMwZip8x78RAyN6ee9NK42WBbKb).

### Annotating training data

We obtained training data from two sources. First, Kyriacou et al. (Kyriacou et al., 2017) manually annotated 52,417 *D. melanogaster* pulse song events from songs that we had recorded previously (Stern, 2014). We downloaded their manual annotations from https://doi.org/10.5061/dryad.80c1f.

Second, before we had completed the *SongExplorer* interface, we employed *Tempo* (https://github.com/JaneliaSciComp/tempo) to manually annotate additional song events. *Tempo* allows simultaneous examination of synced audio and video recordings and manual annotation of song recordings with user-defined song types. We generated dense annotation of 6,770 pulse events of new *D. melanogaster* recordings for the congruence assays. We also generated sparse annotations of 589 sine song events and 172 copulation events from the same *D. melanogaster* recordings annotated by Kyriacou et al. We also generated 15,192 sparse pulse song annotations and 7,537 sparse sine song annotation from five additional strains of *D. melanogaster*. Finally, we annotated 74,065 song events from recordings of *D. erecta, D. mauritiana, D. persimilis, D. pseudoobscura, D. santomea, D. simulans, D. teissieri, D. willistoni* and *D. yakuba*.

Sine song was annotated as a range of time points and pulse events were marked by a single time point per pulse. Pulse annotations were automatically aligned to the largest peak in the full-wave rectified waveform within 5 ms of the manual annotation to reduce variability in annotator pulse placement. Time periods between pulse events that were within the same pulse train were automatically given the annotation of inter-pulse interval (IPI) if the interval was within a median absolute deviation of the species-specific IPI.

### Detecting sound events with time and frequency domain thresholds

Sound events were detected using time domain and frequency domain criteria as follows. Time points with absolute magnitude exceeding a time-domain threshold of 6 median absolute deviations above the median were selected, and gaps between selected time points shorter than 6.4 ms were morphologically closed. A second set of sound events were detected using a frequency-domain threshold of p<0.1 (the FFT window as 25.6 ms and multi-taper settings were NW=4, K=8). Intervals shorter than 25.6 ms were morphologically opened and gaps shorter than 25.6 ms were morphologically closed.

### Recordings of *Drosophila* song

To explore the ability of the deep learning framework to classify diverse song events we recorded male courtship song from seventeen *Drosophila* species using a previously described *Drosophila* courtship song recording apparatus (Arthur et al., 2013). The following fly stocks were employed and samples from the *Drosophila* Species Stock Center (http://blogs.cornell.edu/drosophila/) and Ehime stock center (https://kyotofly.kit.jp/cgi-bin/ehime/) are indicated with the prefixes DSSC and EH: *D. albostrigata* (Kandy, SriLanka, 15112-1811.08); *D. bilimbata* (Guam Island, DSSC 15112-1821.08); *D. elegans* (DSSC 14027-0461.03); *D. erecta* (DSSC 14021-0224.01), *D. gunungcola* (gift of Jonathan Massey); *D. kepulauana* (Sarawak, Borneo Island, DSSC 15112-1761.01); *D. kohkoa* (Sarawak, Borneo Island, DSSC 1511-1771.01); *D. mauritiana* (DSSC 14021-0241.01); *D. melanogaster* (Canton-S); *D. nasuta* (Mombasa, Kenya, DSSC 151121781.06); *D. niveifrons* (Lae, PapuaNewGuinea, EH LAE-221); *D. persimilis* (DSSC 14011-0111.50); *D. pseudoobscura* (DSSC 14022-0121.94); *D. pulau* (Sarawak, Borneo Island, 15112-1801.00); *D. santomea* (STO-CAGO 1482); *D. simulans* (*sim5*, gift of Peter Andolfatto); *D. sulfurigaster* (Kavieng, New Ireland, DSSC 15112-1831.01); *D. taxonf* (Sarawak, Borneo Island, EH B-208); *D. teissieri* (DSSC 14021-0257.01); *D. willistoni* (DSSC 14030-0791.00); and *D. yakuba* (DSSC 14021-0261-01).

### Design and training of the deep neural network classifier

The *SongExplorer* interface enables users to customize the classifier architecture and hyperparameter values to their needs. The classifier tested in this paper is a deep 1D time-domain convolutional neural network implemented in Keras and Tensorflow (https://www.tensorflow.org/) that operates directly on the 5000 Hz waveform. The architecture consists of four convolutional layers with ReLU activation functions. Dropout layers (dropout probability=0.5) were added following each convolution and activation layer. The number of outputs per time point in the output layer corresponded to the number of labels. Eight feature maps were used for the three-word pulse classifier models, and 128 for the others. The last three convolutional layers have a stride of two, which has the effect of reducing the temporal sampling rate of the output predictions by 8-fold. The classifier is trained with cross entropy loss using the Adam optimizer (Kingma and Ba, 2014) with a batch size of 32 and learning rate of 1-e6 for a million training steps over half a day. The learning rate was set such that accuracy on the validation data set had not plateaued until after at least a full epoch of the training data had been sampled. Eight-fold cross-validation of the smallest model (batch=32 features=8) can be done simultaneously on a single Nvidia 1080Ti GPU with only a 29% slow down compared to training a single model. Using seven CPU cores instead of a GPU is only 6% slower for a single model.

### Training and valuating D. melanogaster pulse classifier accuracy and inter-annotator variability with dense annotation

The *D. melanogaster* pulse classifier was trained using 20 densely annotated recordings from Kyriacou et al. (Kyriacou et al., 2017), and the remaining 5 recordings were withheld for validation and used to estimate the best threshold for the classifier probabilities. Each time point was originally annotated with two classes, “pulse” vs “not-pulse”. We added an additional automatically defined class “other pulse” which were pulse-like events originally labeled as “not-pulse”. Events originally labeled “not-pulse” which passed the time domain sound detection threshold but not the frequency domain threshold were given the label of “other pulse”.

An interval of 2 ms was defined around each ground truth pulse. Probabilities predicted by the pulse classifier were thresholded and any contiguous time interval of pulse predictions which overlapped with the 2 ms interval around each ground truth pulse constituted a correct detection or a hit. Predicted pulse intervals that did not overlap with any ground truth intervals constituted false positives, and ground truth intervals that did not overlap any predicted intervals were counted as false negatives or misses. Inter-annotator differences were calculated similarly by defining 2 ms intervals around each annotated pulse and computing overlaps between intervals annotated by different humans. Overlapping intervals were counted as annotations agreed upon by both humans. Unmatched annotations were then labeled “only Person1” or “only Person2.” Precision-recall curves were calculated for both the validation and test set by sweeping the value at which the pulse probability was thresholded. Accuracies were reported for the densely annotated test dataset using threshold values of equal precision and recall on the densely annotated validation set.

### Training and evaluating multi-species song classification from sparse annotations

Multi-species song classification was performed by assigning each time point to the class with the maximum predicted probability. With sparsely annotated ground truth, only annotated time points were considered when evaluating the accuracy of the multi-species classifier. For pulse annotations, a 2 ms window around each pulse was excluded from consideration. The middle half of the interval between two adjacent pulses was automatically annotated with the “IPI” label. For example, if two pulses were annotated at 100 ms and 200 ms, the interval from 125 ms to 175 ms was assigned to the “IPI” class. The multi-species multi-song confusion matrix (Figure 3C) was calculated by counting the number (or percentage) of test dataset time points for which the class with the maximum predicted probability coincided with a ground truth class annotation. Confusion matrices for subsets of the classes (as in Figure 3I,L) were made by first re-normalizing the predicted class probabilities such that the probabilities of the subset of classes sum to 1, and then calculating the confusion matrix for those classes. Precision and recall values for each class in Figure 3D were calculated based on the maximum probability class assignments for each time point, rather than a fixed threshold as used for the pulse classifier.

### Training the species classifier using automatically generated annotations for unsupervised discovery

Sound events from recordings of each species were detected using both time and frequency domain thresholds as described above, and all time points were labeled with the species class label. After training, the classifier was applied to the same detected sounds and the resulting hidden layer activations were used to generate low-dimensional embeddings for interactive discovery of song types.

### Exploratory analysis and visualization using UMAP and t-SNE

*SongExplorer* can perform nonlinear dimensionality reduction of the high-dimensional acoustic signal for visualization in 2D or 3D using the UMAP (McInnes et al., 2018) and t-SNE (van der Maaten and Hinton, 2008) algorithms. Such visualizations can be generated from either the raw acoustic time series representation, a spectrogram representation, or from the intermediate feature representations of a trained deep network.

UMAP was implemented using https://github.com/lmcinnes/umap without any pre-processing of the high dimensional representation. t-SNE was implemented using the *scikit-learn* library, (https://scikit-learn.org/) and utilizes a user-selected initial linear dimensionality reduction using Principle Components Analysis (PCA) to increase algorithm efficiency.

Dimensionality reduction was applied to feature vectors corresponding to 204.8 ms windows of time. For the input layer, the feature vector corresponded to the raw waveform. For all hidden layers, the feature vector was constructed by concatenating the 1D time series of all the feature maps in the hidden layer. The dimensionality of this feature vector is thus given by the number of elements in each hidden layer, given in Figure S1.

## Acknowledgements

We are grateful to Erik Snapp for supervising the Janelia-Loudoun County Public Schools Summer Program, which provided the initial opportunity for co-authors MS and FK to work on this project.

---

**Figure S1.**
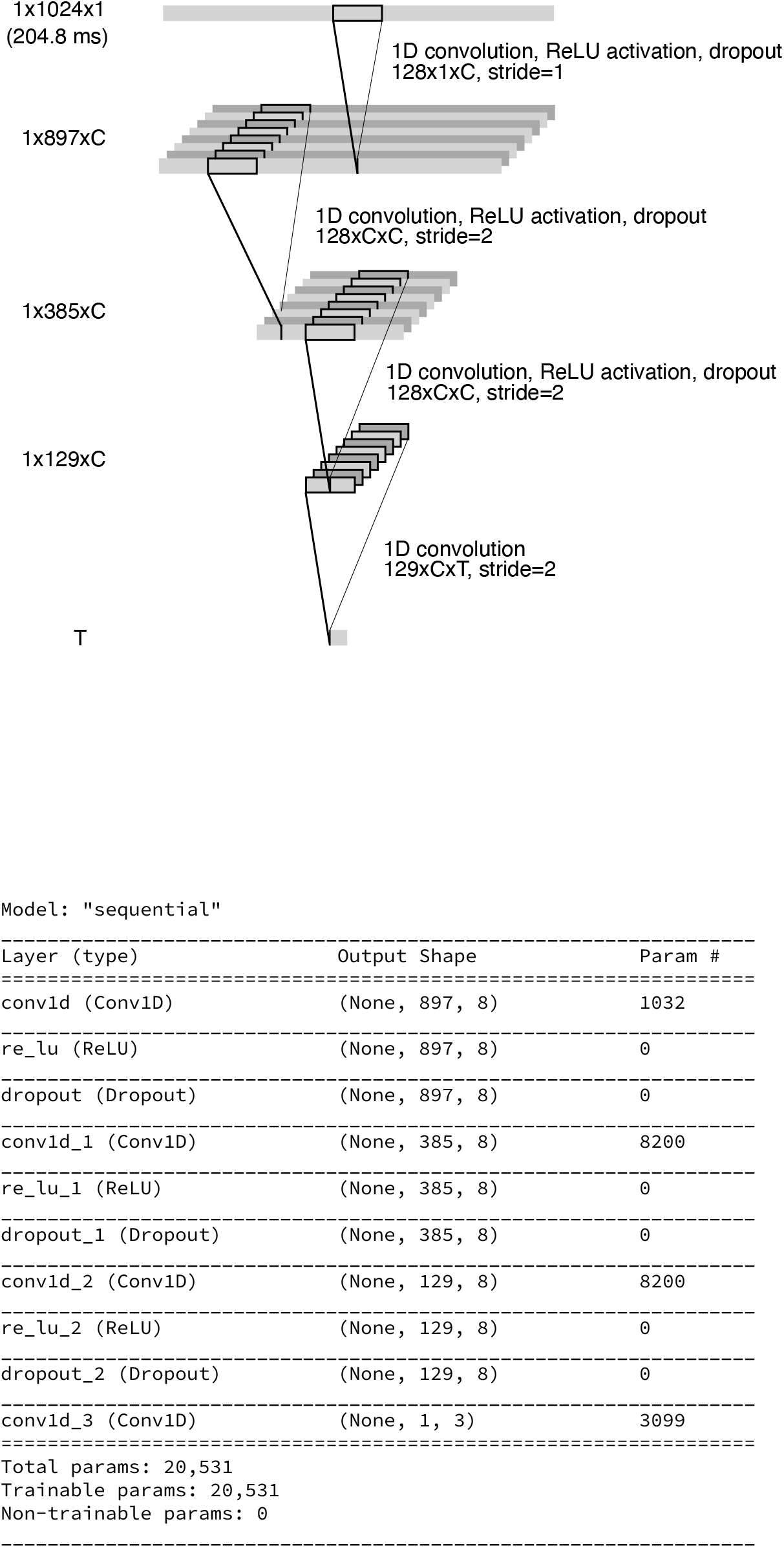
Neural network architecture implemented in *SongExplorer*. Variable C indicates the number of feature maps used in each layer, and was 8 for the 3-word models and 128 for the others. Variable T indicates the number of output classes, which is equal to the number of labelled word classes. Other hyperparameters include a batch size of 32, the Adam optimizer, a dropout probability of 0.5, and a learning rate of 1e-6. A Keras summary of the model is shown below.

**Figure S2.**
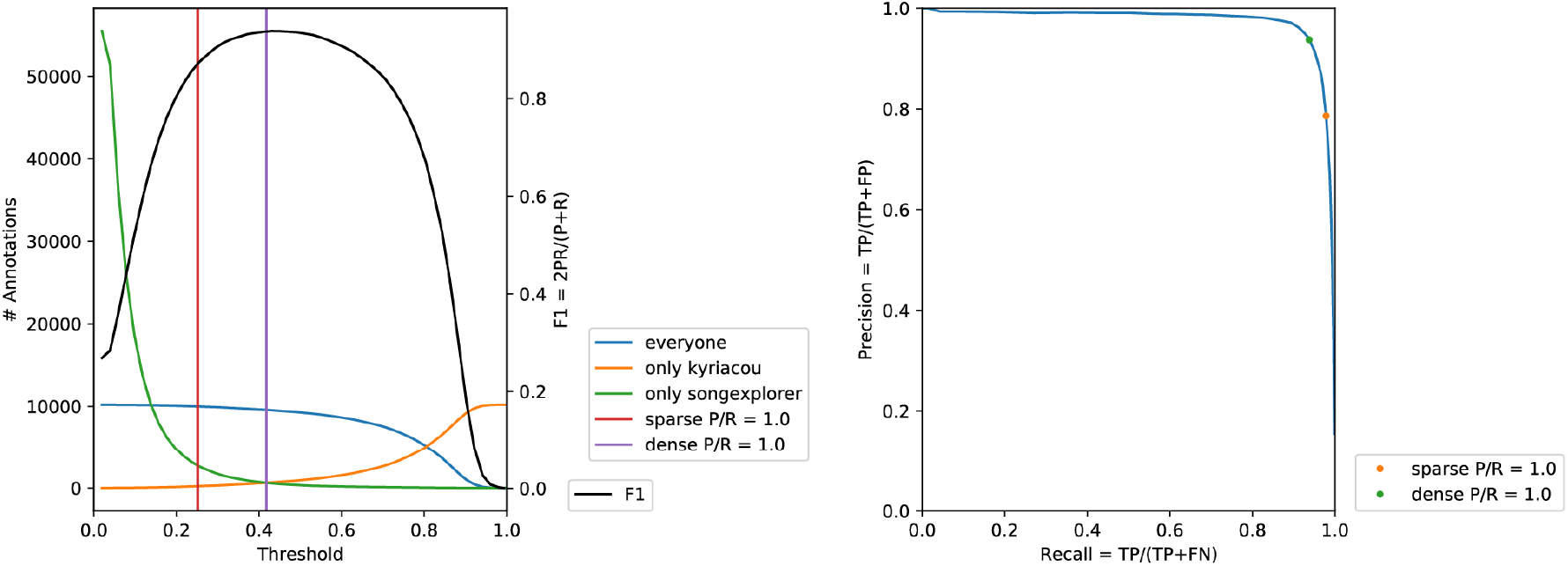
Use of precision-recall curves to select thresholds for converting event probabilities to ethograms. (A) Congruence between dense predictions by SongExplorer and dense annotations by a human was calculated for a range of thresholds. The chosen threshold (vertical purple line) was that for which an equal number of false negatives (orange line) and false positives (green line) was achieved, or whatever the user specified as the desired ratio. Compare with the threshold based on sparse predictions (vertical red line) at just those points which were annotated. (B) Same data as in (A) but plotted parametrically in threshold. The area under the curve is 0.97.

**Figure S3.**
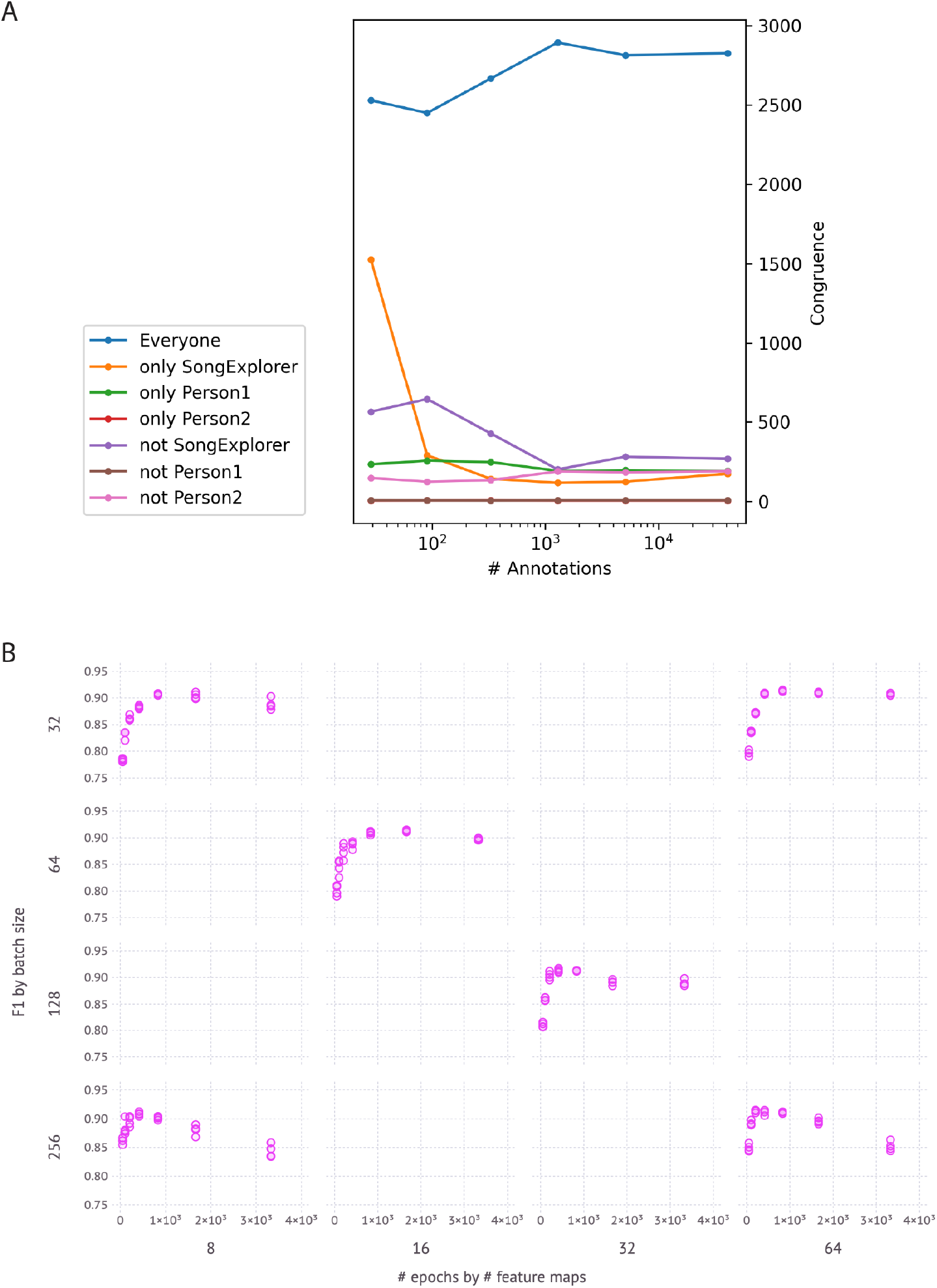
Effect of deep-learning network parameters on trained model accuracy. Maximum F1 is relatively insensitive to 8-fold changes in batch size and the number of feature maps. Larger batch sizes are more susceptible to overtraining, and fewer feature maps require longer training times and exhibit larger variability across repetitions. The learning rate was set such that accuracy on the validation data set had not plateaued until after at least a full epoch of the training data had been sampled (1e-6 for all models except 1e-7 for batch=32 features=64 and 1e-5 for batch=256 features=8). Eight-fold cross-validation of the smallest model (batch=32 features=8) can be done simultaneously on a single Nvidia 1080Ti GPU with only a 29% slow down compared to training a single model. Using seven CPU cores instead of a GPU is only 6% slower for a single model.

**Figure S4.**
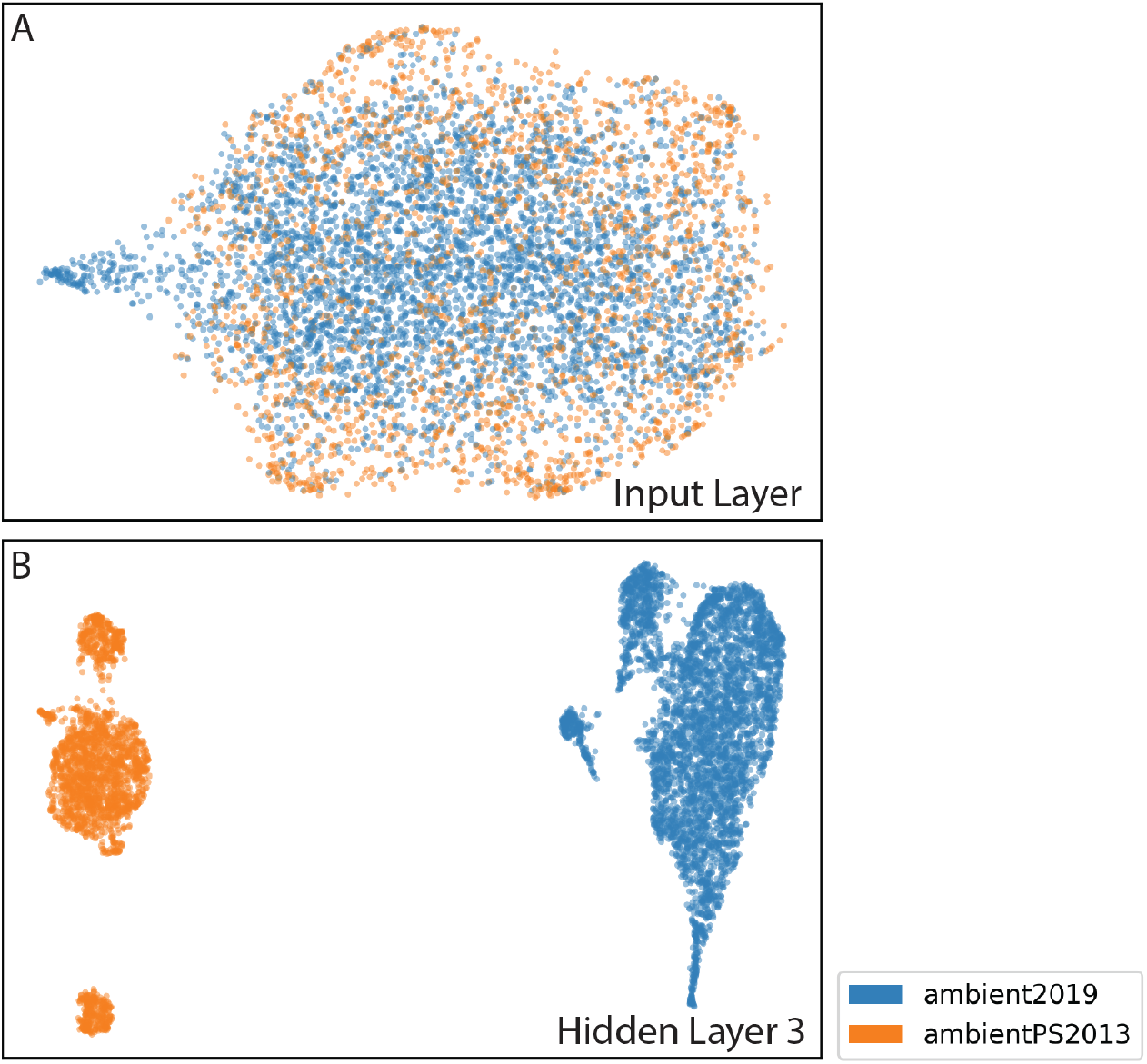
Recordings in different years display different noise characteristics. Recordings performed in 2013 and 2019 were made using chambers of different sizes. We tested whether recordings made in 2013 and 2019 differed systematically in their noise characteristics by training the deep learning network to recognize ambient sounds in recordings from these two years. UMAP projections of the ambient noise from each year mostly overlap in the input layer (A), but are strongly differentiated in the third hidden layer (B), implying that the classifier differentiated the two song types solely on the basis of ambient noise.

**Figure S5.**
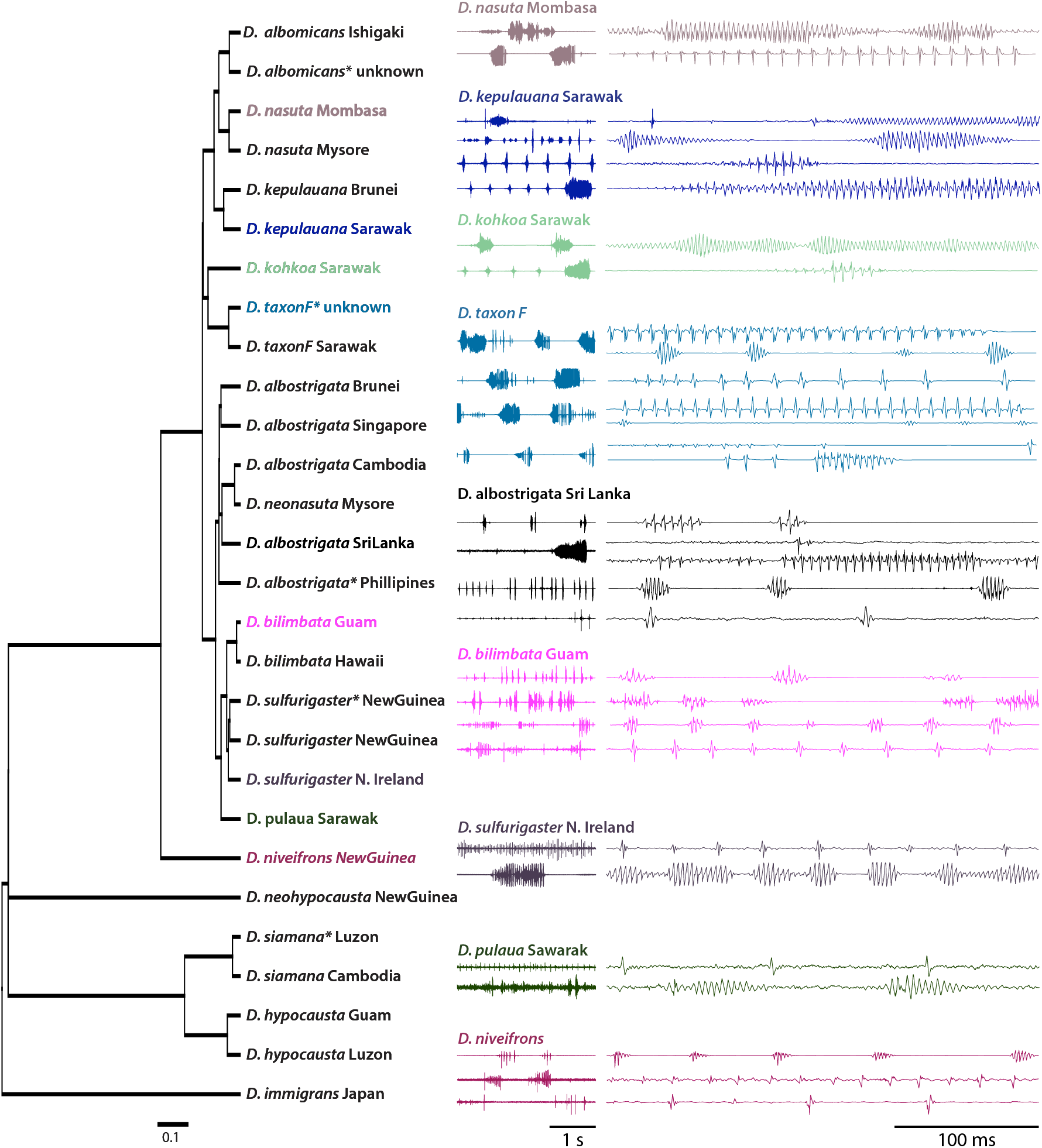
Dependence of classifier performance on the number of labelled song events used for training. As few as 100 labelled song events resulted in performance approximately as high as songs events labelled with many more events.

**Figure S6.** Song types discovered for nine species of the *D. nasuta* species group. Phylogeny of the species examined is shown on the left, with the samples used for song analysis in the same color font as the songs shown on the right. One previous study had found one song type each for 6 of the species we studied and none for two of the species we studied. In contrast, we identified song in all species we studied, and from two to seven apparently distinct song types in different species.

Figure S7 Video. Videos illustrating the *SongExplorer* workflow can be found at the following YouTube channel: https://www.youtube.com/playlist?list=PLYXyXDkMwZip8x78RAyN6ee9NK42WBbKb

Supplementary Audio Files: WAV files of the song types illustrated in Figure S5 are available on FigShare.

